# An integrated approach to identifying sex-specific genes, transcription factors, and pathways relevant to Alzheimer’s disease

**DOI:** 10.1101/2023.09.05.556293

**Authors:** Adolfo López-Cerdán, Zoraida Andreu, Marta R. Hidalgo, Irene Soler-Sáez, Antonio Porlan, Macarena Pozo-Morales, Santiago Leon, María de la Iglesia-Vayá, Akiko Mikozami, Franca R. Guerini, Francisco García-García

## Abstract

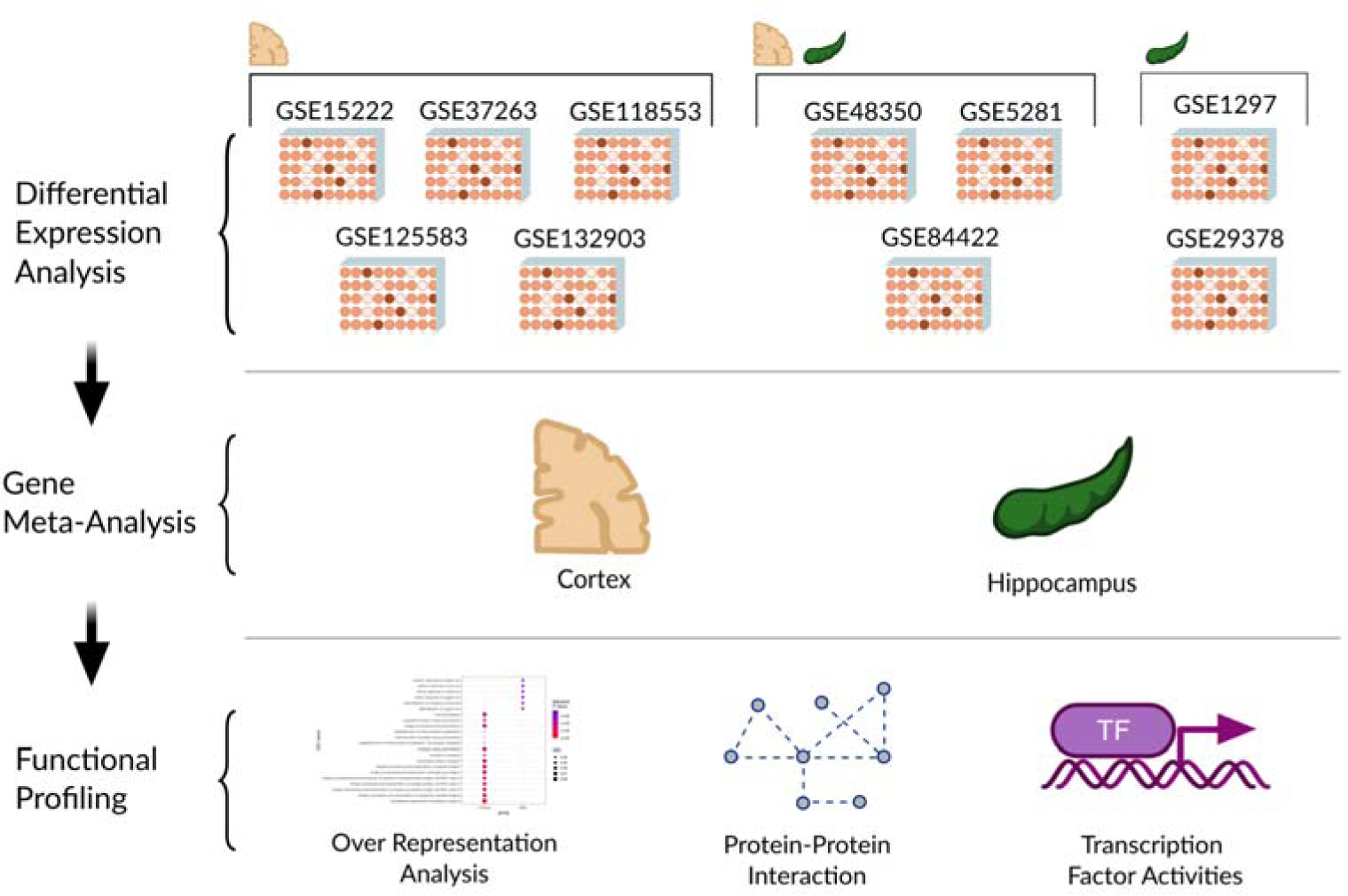

**Background:** Age represents a significant risk factor for the development of Alzheimer’s disease (AD); however, recent research has documented an influencing role of sex in several features of AD. Understanding the impact of sex on specific molecular mechanisms associated with AD remains a critical challenge to creating tailored therapeutic interventions.

**Methods:** The exploration of the sex-based differential impact on disease (SDID) in AD used a systematic review to first select transcriptomic studies of AD with data regarding sex in the period covering 2002 to 2021 with a focus on the primary brain regions affected by AD - the cortex (CT) and the hippocampus (HP). A differential expression analysis for each study and two tissue-specific meta-analyses were then performed. Focusing on the CT due to the presence of significant SDID-related alterations, a comprehensive functional characterization was conducted: protein-protein network interaction and over-representation analyses to explore biological processes and pathways and a VIPER analysis to estimate transcription factor activity.

**Results:** We selected 8 CT and 5 HP studies from the Gene Expression Omnibus (GEO) repository for tissue-specific meta-analyses. We detected 389 significantly altered genes in the SDID comparison in the CT. Generally, female AD patients displayed more affected genes than males; we grouped said genes into six subsets according to their expression profile in female and male AD patients. Only subset I (repressed genes in female AD patients) displayed significant results during functional profiling. Female AD patients demonstrated more significant impairments in biological processes related to the regulation and organization of synapsis and pathways linked to neurotransmitters (glutamate and GABA) and protein folding, Aβ aggregation, and accumulation compared to male AD patients. These findings could partly explain why we observe more pronounced cognitive decline in female AD patients. Finally, we detected 23 transcription factors with different activation patterns according to sex, with some associated with AD for the first time. All results generated during this study are readily available through an open web resource Metafun-AD (https://bioinfo.cipf.es/metafun-ad/).

**Conclusion:** Our meta-analyses indicate the existence of differences in AD-related mechanisms in female and male patients. These sex-based differences will represent the basis for new hypotheses and could significantly impact precision medicine and improve diagnosis and clinical outcomes in AD patients.

**Highlights:**

- Female AD patients possess more affected genes than male AD patients.
- 389 genes from the sex-based differential impact on disease comparison significantly impact the cerebral cortex and suggest a more significant effect on cognitive function in female AD patients.
- The cluster of repressed genes in female AD patients functionally impacts glutamate and GABA neurotransmitters and Aβ deposition.
- Female AD patients exhibit several transcription factors with significantly different activity patterns compared to male AD patients.
- This work includes Metafun-AD, an open and interactive web tool to explore all generated data and results.

## BACKGROUND

AD represents the most common form of dementia in the aged population. The latest update from the World Health Organization (WHO, 2023) indicates that more than 55 million persons have dementia worldwide, with AD representing 60-70% of the cases. Of note, AD incidence is rising fast (about 10 million new cases each year), with over 131 million AD cases estimated to occur by 2050 [1]. Currently, AD remains incurable and terminal and represents a leading cause of dependency, disability, and mortality [2, 3].

The brain of AD patients suffers from significant levels of neuronal death. The neuropathology relates to the accumulation of amyloid-beta (Aβ) protein plaques and neurofibrillary tangles of the Tau protein [4, 5]. Excess Aβ plaques and Tau tangles prompt processes disrupting neuronal communication, metabolism, and repair, thereby disrupting homeostasis [6]. The limbic system (and the HP in particular, which remains critical to the formation of new memories and learning) represents the first point of attack. Subsequently, AD sufferers undergo deterioration in the CT, resulting in the inability to control emotional outbursts and carry out daily tasks. Finally, the brainstem becomes damaged in advanced stages, which causes organ failure. In general, AD is clinically characterized by memory loss but presents with other cognitive and behavior-related symptoms.

The causes of AD remain relatively unknown. While genetic factors seem to determine early-onset AD, the interaction between risk and environmental factors may drive late-onset AD, which encompasses 95% of all cases [7]. Several studies have also highlighted sex as a critical risk factor in neurodegenerative diseases and the development of late-onset AD [8–11]. Females comprise two-thirds of all AD patients [12] and sex-related differences are evident in patterns of disease manifestation and the rates of cognitive decline and brain atrophy, suggesting sex as a crucial variable in disease heterogeneity [13–19]. Nevertheless, we understand relatively little about the molecular mechanisms for the evident sex bias in AD patients; however, multi-omics analyses and datasets from human AD samples and animal models offer an excellent platform to study sex-related molecular and pathway alterations. Including sex as a variable in AD research will improve precision medicine strategies and provide for more rapid advances in diagnosis and treatment.

To investigate sex-based differences in AD, we provide two independent meta-analyses of gene expression datasets that consider the sex of human AD patients. We performed one meta-analysis in the CT (eight studies) and the other in the HP (five studies), given that the CT and HP represent the two central brain regions affected by AD pathogenesis. Our *in-silico* approach revealed the expression of differentially-expressed gene subsets in female and male AD patients. At a functional level, these gene clusters mainly involve functions related to neurotransmission regulation, synapses and pre-synapses, calcium signaling, pH reduction, MAPK activity, ubiquitination, and protein folding. Additionally, female AD patients presented with the significantly upregulated expression of transcription factors with different activity patterns (ADNP, HMGN3, IRF3, KLF5, KLF9, MAZ, MBD3, MYNN, PRDM14, SIX5, and ZNF207 – activated; GTF2B, HOXB13, NANOG, NME2, PCGF2, SNAI2, ZBTB7A, ZC3H8, ZHX1, and ZHX2 - repressed). Additionally, two transcription factors (CEBPZ and TERFS) displayed divergent expression patterns by sex. We highlight the freely available nature of our results in the Metafun-AD web tool as a starting point for future studies.

## METHODS

All bioinformatics and statistical analysis were performed using R version 4.2.1 software [20].

### Study Search and Selection

Available datasets were collected from the Gene Expression Omnibus (GEO) [21] public repository. A systematic search of all published studies in public repositories (2002-2021) was conducted during 2021, following the preferred reporting items for systematic reviews and meta-analyses (PRISMA) guidelines [22]. Keywords employed in the search were “Alzheimer,” “Alzheimer’s Disease”, and “AD”. The following inclusion criteria were applied:

● Transcriptomic studies on *Homo sapiens*
● Control and AD-affected patients included
● Sex, disease/control status, age, and brain region variables registered
● RNA extracted directly from post-mortem brain tissues (no cell lines or cultures)
● Brain tissues from either the CT or HP
● Sample size > 3 for case and control groups in both sexes

Normalized gene expression data of 8 microarray AD datasets (GSE118553, GSE1297, GSE132903, GSE15222, GSE29378, GSE37263, GSE48350, GSE5281, and GSE84422) and the raw counts’ matrix of the GSE125583 RNA-sequencing study from the GEO repository were retrieved. Data downloading was performed using the GEOquery R package [23] for microarray studies and manually for the RNA-sequencing study.

### Individual Transcriptomics Analysis

For each selected study, an individual transcriptomics analysis comprised two steps: preprocessing and differential expression analysis.

Data preprocessing included the standardization of the terminology for the clinical variables in each study, the homogenization of gene annotation, and exploratory data analysis. For the microarray datasets, the normalization methods performed by the original authors were assessed, log_2_ transforming data matrices when necessary. All probe sets were annotated to HUGO gene symbols [24] using the annotation provided by each microarray platform. The median of expression values was calculated when dealing with duplicated probe-to-symbol mappings. For the RNA-sequencing dataset, the count matrix was preprocessed using the edgeR R package [25] and transformed using the *Voom* function included in the limma R package [26]. The exploratory analysis included unsupervised clustering and PCA to detect expression patterns between samples and genes and the presence of batch effects in each study.

Differential expression analyses were performed using the limma R package to detect the sex-based differential expression of genes. To achieve this goal, the following comparison was applied:

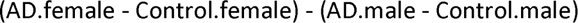

This comparison allows the detection of genes with a sex-based differential impact on disease (SDID). Genes with a log_2_ fold change (LFC) greater than zero show either a higher increase or a lesser decrease in expression in female AD patients when comparing the effect of the disease between sexes. On the contrary, genes with an LFC lower than zero have a higher increase or a lesser decrease in expression in male AD patients when comparing the effect of the disease between sexes.

To gain a better understanding of the sex-based differential behavior detected with the previous comparison, two additional comparisons were performed: A case vs. control comparison performed only in females (AD.female - Control.female) that evaluates the impact of AD in females (IDF) and another performed only in males (AD.male - Control.male), that informs us about the impact AD in males (IDM).

These comparisons were applied separately to all CT and HP samples. The patients’ age was included in the limma linear model as a blocking factor to reduce its impact on the results. P-values were corrected using the Benjamini-Hochberg procedure [27] and considered significant when below a threshold of 0.05.

### Gene Expression Meta-Analysis

Differential gene expression results were integrated into a meta-analysis [28] for each brain region (CT and HP). Meta-analyses were implemented with the R package metafor [29] under the DerSimonian & Laird random-effects model [30], considering individual study heterogeneity. This model considers the variability of individual studies by increasing the weights of studies with less variability when computing meta-analysis results. Thus, the most robust functions between studies are highlighted.

P-values, corrected p-values, LFC, LFC standard error (SE), and LFC 95% confidence intervals (CI) were calculated for each evaluated gene. Genes with corrected p-values lower than 0.05 were considered significant, and both funnel and forest plots were computed for each. These representations were evaluated to assess for possible biased results, where the LFC represents the effect size of a gene, and the SE of the LFC serves as a study precision measure [31]. Sensitivity analysis (leave-one-out cross-validation) was conducted for each significant gene to verify alterations in the results due to the inclusion of any study. The Open Targets platform (release 22.09) [32] was used to explore the associations of significant genes with AD.

### Sex-based Functional Signature in the CT

Gene meta-analysis of CT data revealed gene sets with a significantly differential expression pattern between male and female AD patients. Several analyses were conducted to identify the functional implications of these differences.

Over-Representation Analysis (ORA) [33] through clusterProfiler and ReactomePA R packages [34, 35] was first used to determine the biological functions and pathways overrepresented in all significant gene subsets. P-values and corrected p-values were calculated for each GO (Gene Ontology) term from the “Biological Processes” GO ontology [36] and each Reactome pathway [37]. Every function and pathway with a corrected p-value lower than 0.05 was labeled as over-represented in each gene set.

Protein-protein interaction (PPI) networks were then calculated using the STRING web tool for each subset of genes [38]. The total number of edges was examined, and PPI enrichment was assessed using the default parameters for each network.

A VIPER analysis [39] with human regulons obtained from the DoRothEA R package [40] was performed to estimate transcription factor activity. Regulons with a confidence level of A, B, C, or D were selected, excluding those with less than 25 genes (n = 217). The p-values were corrected using the Benjamini & Hochberg method. Normalized enrichment scores (NES) were calculated by VIPER as a measure of relative transcription factor activity.

### Metafun-AD Web Tool

All data and results generated in the different steps of the meta-analysis are available in the Metafun-AD web tool [41], which is freely accessible to any user and allows the confirmation of the results described in this manuscript and the exploration of other results of interest. The front end was developed using Quarto 1.2 [42], and the interactive graphics used in this web resource have been implemented with plotly [43].

This easy-to-use resource is divided into eight sections: 1) framework and summary of analysis results in each phase. Then, 2) systematic review conducted to identify studies and, for each of them, the detailed results of the 3) exploratory analysis and differential expression. 4) The gene meta-analysis results from the different meta-analyses. Sections 5-7) provide detailed tables and figures corresponding to the results of the three functional profiling methods (ORA, PPI, and Transcription Factor Activity). Finally, section 8) provides a synthesis of the bioinformatics methods. Through the web, the user can interact with the web tool through graphics and tables and search for specific information for a gene or function.

## RESULTS

In this study, we aimed to investigate the existence of sex-based differences associated with AD using a systematic review and two meta-analyses of transcriptomics studies. We obtained data from those studies that included information on the sex of the patients from the GEO repository. We conducted one meta-analysis for each of the two primary brain regions affected by AD pathogenesis - CT (eight studies) and HP (five studies). Subsequently, we explored the biological implications of the CT meta-analysis results by utilizing three distinct functional profiling methods - ORA, PPI network construction, and transcription factor activity analysis.

### Systematic Review and Study Selection

In our search for studies on AD, we identified 76 non-duplicated entries, 40 of which (52%) included both male and female patients. After applying inclusion and exclusion criteria (Methods, **Figure 1**), we selected 14 studies for comparison; however, we excluded 4 after an exploratory analysis. Thus, we analyzed 10 studies comprising 2508 samples (909 controls and 1599 AD cases) from the CT and HP (**Table 1**). **Figure 2** presents the sex distribution by study and brain region, with an overall proportion of 44% males and 56% females. The median age of the participants was 85. **Table 1** and **Figure 2** report additional information on the selected studies and their clinicopathological features.

**Figure 1.**
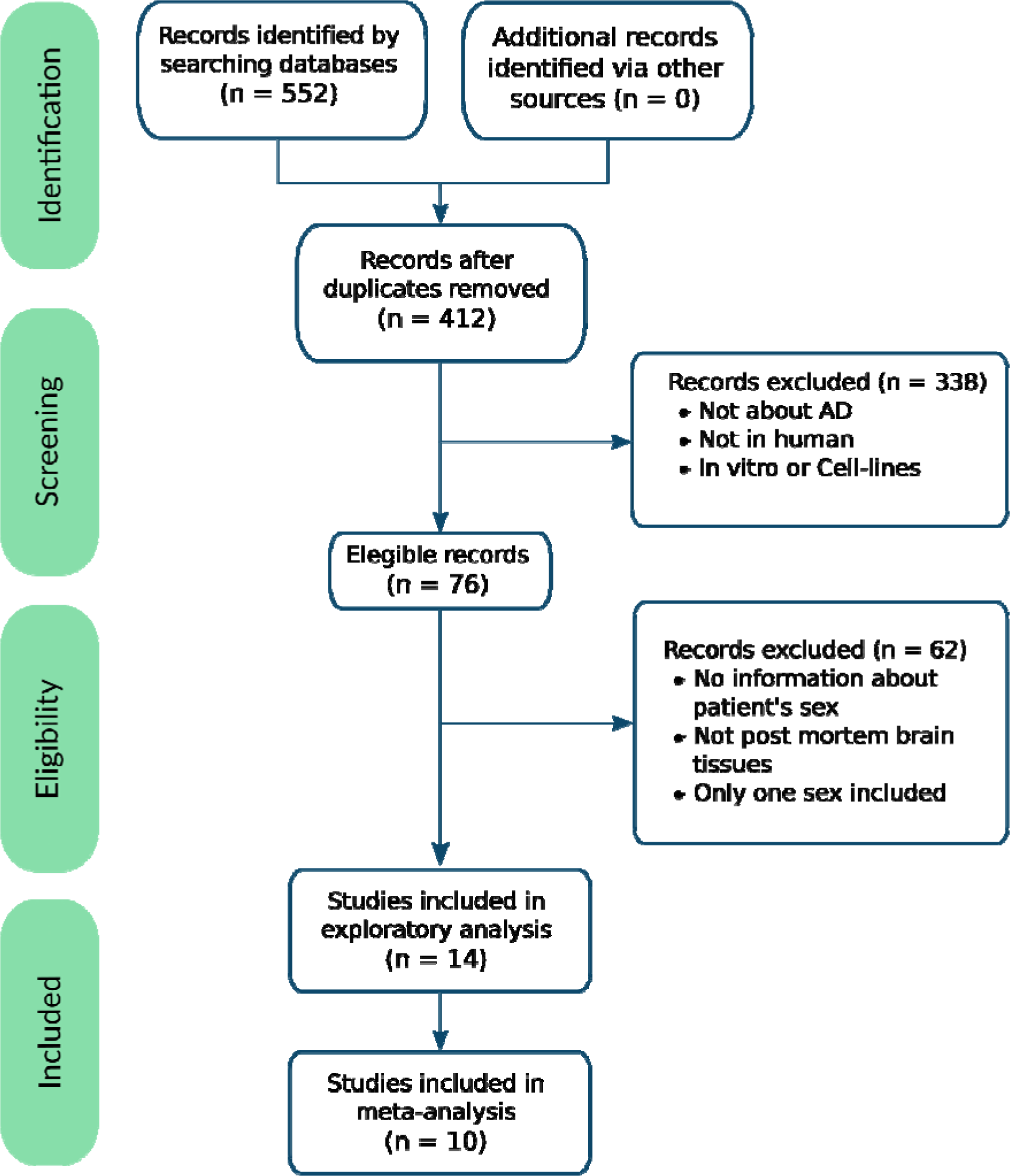
The flow of information through the distinct phases of the systematic review following PRISMA statement guidelines [22].

**Figure 2.**
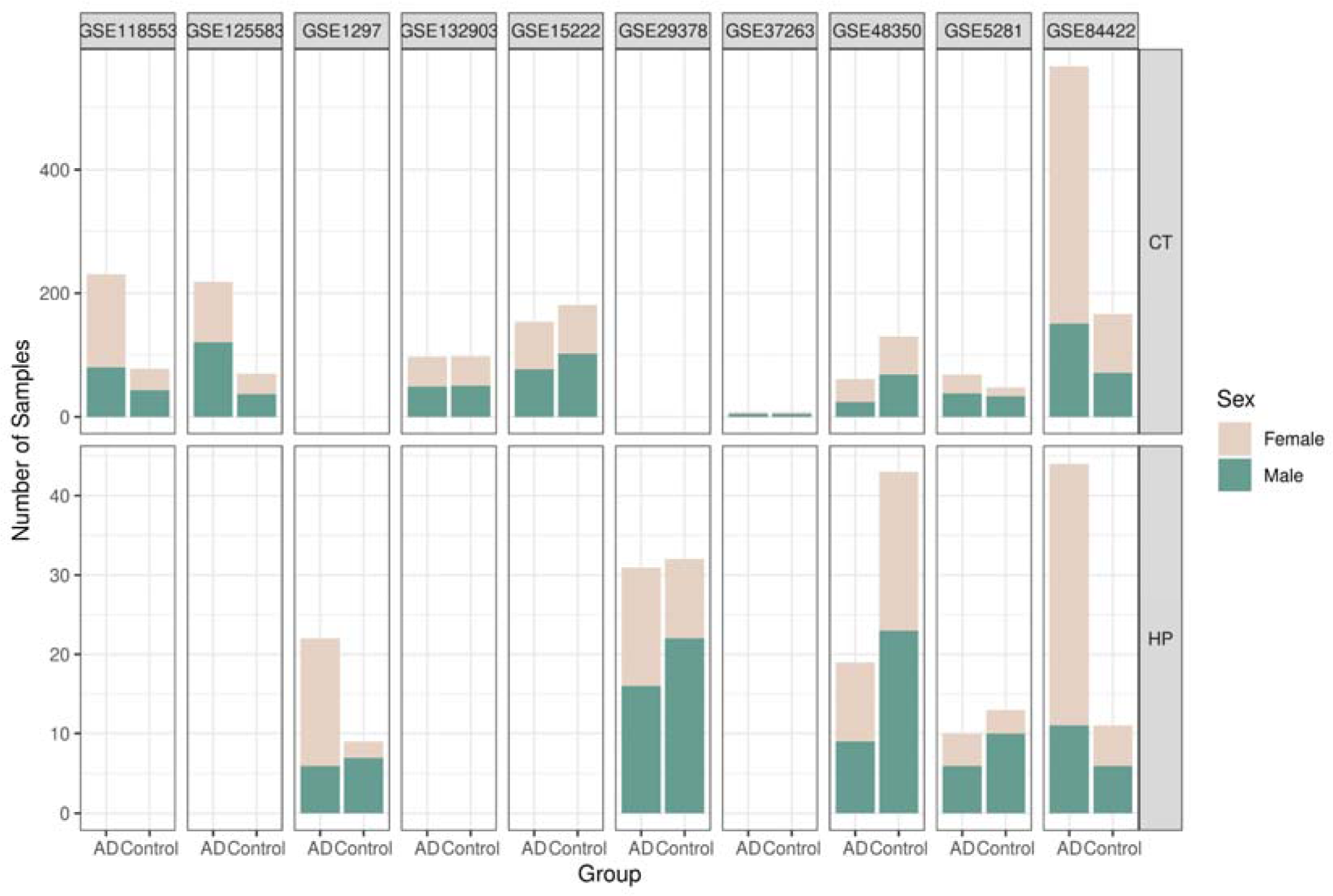
The number of samples per study divided by sex, study, and experimental group. AD – Alzheimer’s disease, CT – cortex, HP – hippocampus.

**Table 1.** Studies selected after the systematic review.

| Study | Platform | Brain area | Publication (PMID) |
| --- | --- | --- | --- |
| GSE118553 | Illumina HumanHT-12 V4.0 expression beadchip | CT | [44] |
| GSE125583 | Illumina HiSeq 2500 | CT | [45] |
| GSE1297 | Affymetrix Human Genome U133A Array | HP | [46] |
| GSE132903 | Illumina HumanHT-12 V4.0 expression beadchip | CT | [47] |
| GSE15222 | Sentrix HumanRef-8 Expression BeadChip | CT | [48] |
| GSE29378 | Illumina HumanHT-12 V3.0 expression beadchip | HP | [49] |
| GSE37263 | Affymetrix Human Exon 1.0 ST Array | CT | [50] |
| GSE48350 | Affymetrix Human Genome U133 Plus 2.0 Array | CT, HP | [51] |
| GSE5281 | Affymetrix Human Genome U133 Plus 2.0 Array | CT, HP | [52] |
| GSE84422 | Affymetrix Human Genome U133A/U133B Arrays | CT, HP | [53] |

Brain areas: Cortex (CT) and Hippocampus (HP)

### Individual Transcriptomic Analysis

We conducted exploratory and processing steps on the datasets to ensure their comparability in subsequent analyses. We applied log_2_ transformation to studies GSE1297, GSE5281, GSE15222, GSE29378, and GSE48350 to homogenize magnitude order and then filtered out samples from regions different from CT and HP from all studies. We excluded studies GSE44768, GSE44770, GSE44771, and GSE33000 from the selection.

The differential expression results for each study provided a variable number of significantly altered genes across comparisons and studies (**Table 2**). In the case of the SDID comparison, only GSE15222, GSE29378, and GSE84422 reported the significantly altered expression of genes.

**Table 2.** Summary of differential gene expression analysis by brain region and comparison.

|  | Significantly Altered Genes: |  |  | Significantly Altered Genes: |  |  | Significantly Altered Genes: |  |  |
| --- | --- | --- | --- | --- | --- | --- | --- | --- | --- |
|  | IDF |  |  | IDM |  |  | SDID |  |  |
|  | LFC > 0 | LFC < 0 | Total | LFC > 0 | LFC < 0 | Total | LFC > 0 | LFC < 0 | Total |
| GSE118553 (CT) | 387 | 291 | 678 | 759 | 401 | 1160 | 0 | 0 | 0 |
| GSE125583 (CT) | 4461 | 4230 | 8691 | 3366 | 3482 | 6948 | 0 | 0 | 0 |
| GSE1297 (HP) | 2 | 0 | 2 | 0 | 0 | 0 | 0 | 0 | 0 |
| GSE132903 (CT) | 5925 | 4782 | 10707 | 4643 | 4473 | 9116 | 0 | 0 | 0 |
| GSE15222 (CT) | 5042 | 3699 | 8741 | 2054 | 1628 | 3682 | 267 | 148 | 415 |
| GSE29378 (HP) | 63 | 21 | 84 | 347 | 193 | 540 | 0 | 1 | 1 |
| GSE37263 (CT) | 0 | 0 | 0 | 0 | 0 | 0 | 0 | 0 | 0 |
| GSE48350 (CT) | 28 | 34 | 62 | 2 | 6 | 8 | 0 | 0 | 0 |
| GSE48350 (HP) | 760 | 421 | 1181 | 15 | 1 | 16 | 0 | 0 | 0 |
| GSE5281 (CT) | 718 | 2307 | 3025 | 618 | 3815 | 4433 | 0 | 0 | 0 |
| GSE5281 (HP) | 96 | 70 | 166 | 138 | 111 | 249 | 0 | 0 | 0 |
| GSE84422 (CT) | 109 | 235 | 344 | 355 | 490 | 845 | 174 | 89 | 263 |
| GSE84422 (HP) | 0 | 0 | 0 | 0 | 0 | 0 | 0 | 0 | 0 |

CT = cortex tissue, HP = hippocampus tissue, IDF = Impact of the disease in females, IDM = Impact of the disease in males, and SDID = sex-based differential impact on disease

### Gene-expression meta-analyses reveal sex differences in the cortex

For each different comparison (IDF, IDM, and SDID), we performed a gene expression meta-analysis in the HP and CT, analyzing eight studies for the CT and five for the HP. We employed the SDID comparison to determine the genes with sex-based differential expression patterns in AD. We then performed the IDF and the IDM comparisons to provide more details regarding the impact of sex in this subset of genes.

The SDID comparison revealed no significantly altered gene expression in the HP; therefore, we excluded this region from further functional analyses. The SDID comparison did, however, detect 389 genes with significantly altered expression in the CT. Additionally, the IDF and IDM comparisons revealed 3763 and 1876 differentially expressed genes in AD in females and males, respectively (**Supplementary Table 1**).

We divided the 389 genes with significantly altered gene expression in the SDID comparison according to their significance and LFC value in the IDF, IDM, and SDID comparisons (**Figures 3A and 3B**). The six resultant subsets of genes were:

1. More repressed genes in AD females (179 genes)
2. More expressed genes in AD females (57 genes)
3. More repressed genes in AD males (14 genes)
4. More expressed genes in AD males (6 genes)
5. Increased in AD females related to AD males (69 genes)
6. Decreased in AD females related to AD males (64 genes)

**Figure 3.**
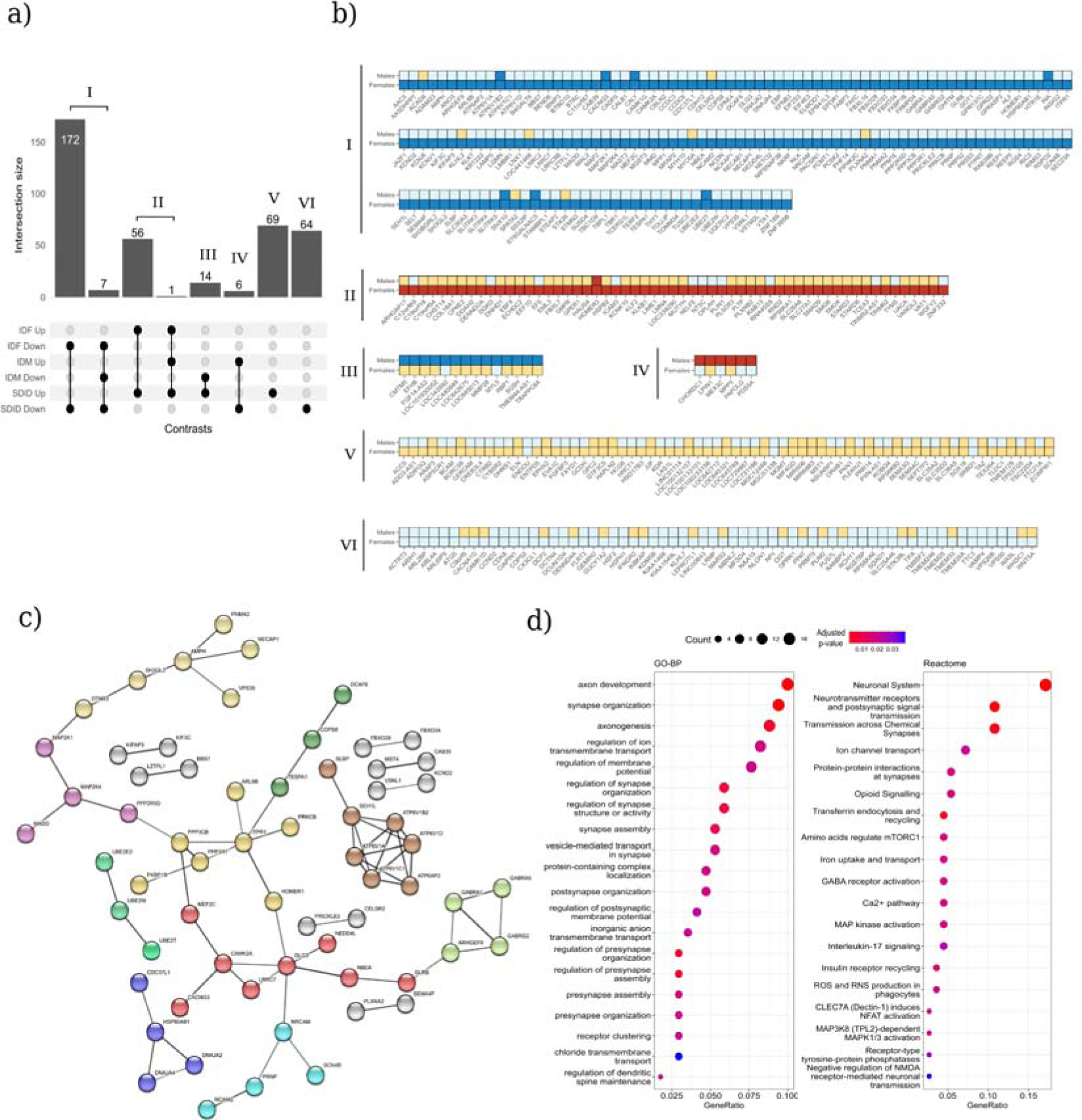
Genes with a sex-based differential impact on disease detected in the cerebral cortex. **A**) Upset plot of the intersections between the differentially expressed genes encountered in the IDF, IFM, and SDID comparisons. Only the intersections containing significantly altered gene expression in the SDID are shown. **B**) Tile plot of every gene composing the six resulting Subsets (I-VI). The direction of expression and significance are reflected for female and male patients, with the following color pattern: blue - underexpression, red - overexpression, and darker colors - statistical significance (adjusted p-value < 0.05). **C**) PPI network calculated from gene Subset I, showing only network edges with an interaction score greater than 0.7. Network nodes are colored according to their cluster (See **Table 3**). **D**) Dot Plots summarizing the significant GO biological processes (left) and Reactome pathways (right) detected by ORA in gene Subset I.

**Table 3.**
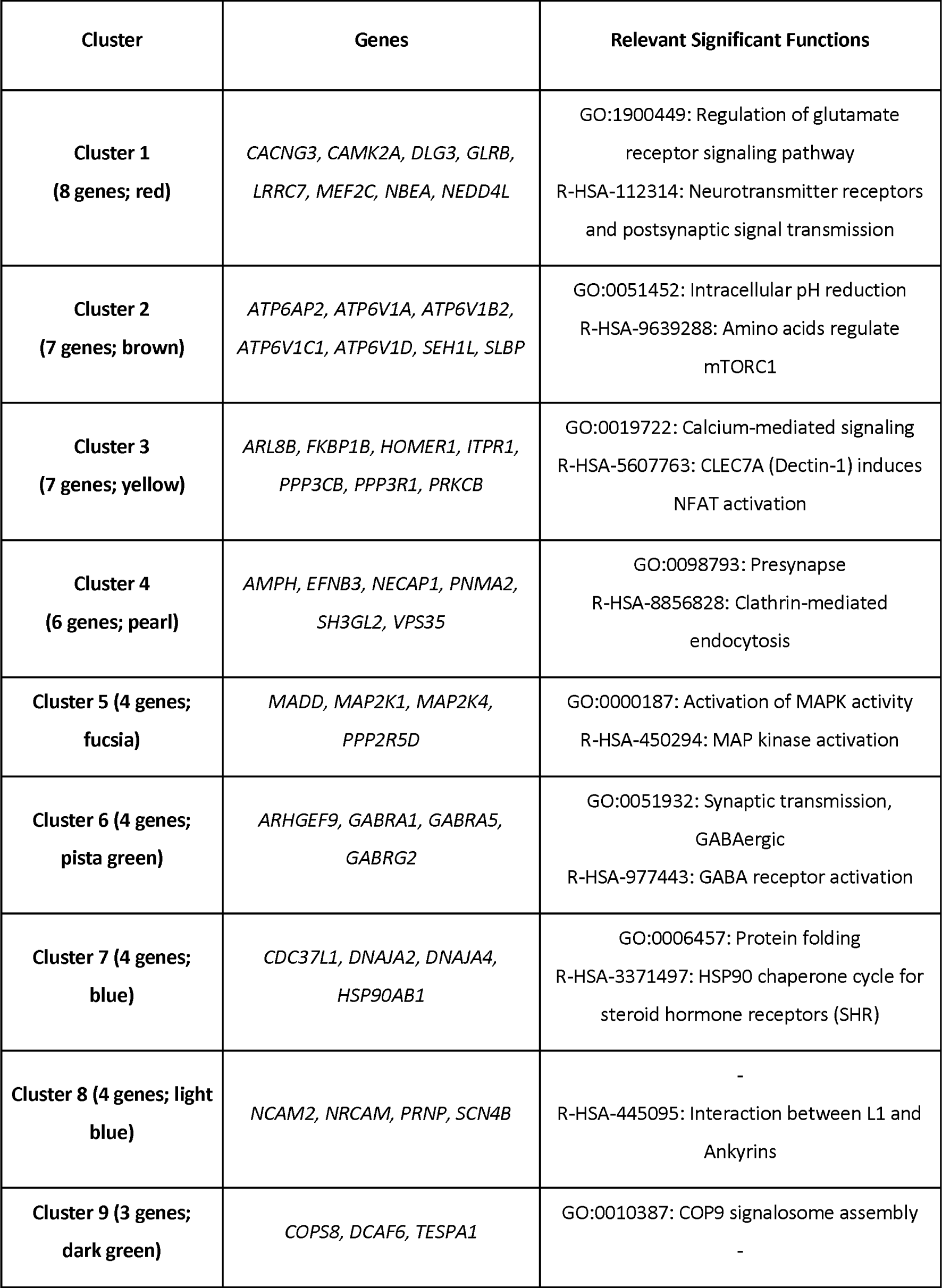
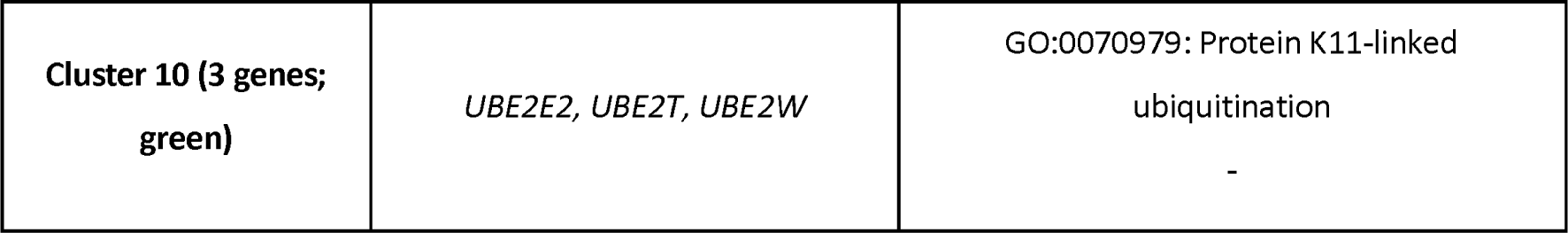
Gene clusters extracted from the PPI network calculated from gene Subset I. *Significantly repressed genes in affected females*

### Functional profiling of significant genes in the CT

We functionally characterized the gene subsets described above through ORA and the calculation of PPI networks; however, we only obtained significant results in the “More repressed genes in AD females” subset (Subset I).

ORA results with gene subset I revealed a significant overrepresentation of 34 GO terms (BP ontology) and 20 Reactome pathways (**Figure 3D**, **Supplementary Table 3**). Most significant GO terms related to processes involved in synaptic organization (GO:0050808, GO:0099174, GO:0050807, GO:0099173), assembly (GO:1905606, GO:0099054, GO:0051963), and vesicle-mediated transport (GO:0099003), and the regulation of membrane potential (GO:0042391, GO:0060078, GO:1904062), axonogenesis (GO:0061564, GO:0007409), and cytoskeleton-dependent intracellular transport (GO:0030705), which play essential roles in the development and function of neurons. Most significant Reactome pathways also participated in the neuronal system (R-HSA-112316), including Neurotransmitter receptors and postsynaptic signal transmission (R-HSA-112314), GABA receptor activation (R-HSA-977443), Ca2+ pathway (R-HSA-4086398), and Negative regulation of NMDA receptor-mediated neuronal transmission (R-HSA-9617324).

The calculated PPI network revealed significantly more interactions between genes than expected for a random set of genes of the same size and degree distribution, with a PPI enrichment p-value of 1.27 x 10-11 (**Figure 3C**). We computed gene clusters using the MCL clustering method included in the STRING web tool. As a result, we elucidated 10 clusters composed of at least 3 genes, most functionally related to neurological processes such as synapsis (**Table 3**). STRING provided the functional enrichments in each cluster (**Supplementary Table 2**).

### Transcription Factors Activity in the CT

The transcription factor activity analysis (VIPER analysis) of the consensus profiles calculated from the three meta-analyses (IDF, IDM, and SDID) reported 23 transcription factors with sex-affected patterns of expression (adjusted p-value < 0.05 in SDID). We split these transcription factors into three groups according to their profile: “Activated,” “Repressed,” and “Divergent” (**Figure 4**).

**Figure 4.**
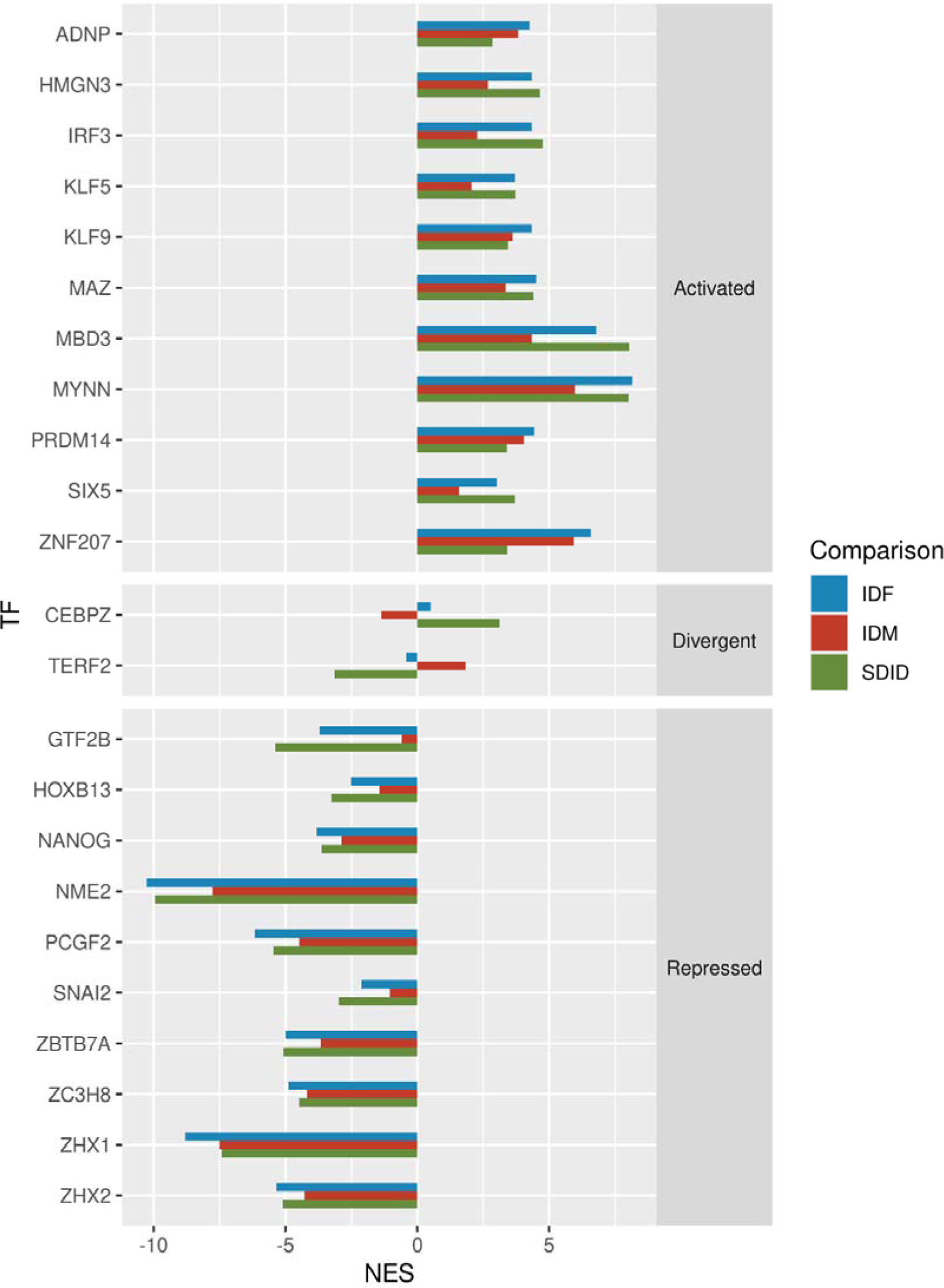
Transcription factors with significantly altered activity calculated from the SDID comparison. (adjusted p-value < 0.05). Activation values are measured as normalized enrichment scores (NES). Transcription factors labeled as “Activated” display positive values of NES in all comparisons, “Repressed” display negative values of NES in all comparisons, and “Divergent” display qualitative differences in their activation across the IDF, IDM, and SDID comparisons.

We labeled as Activated the eleven transcription factors with NES of > 0 in the IDF, IDM, and SDID comparisons. These transcription factors are predicted to be activated in both female and male AD patients but significantly increased in females compared to males (Genes - *ADNP, HMGN3, IRF3, KLF5, KLF9, MAZ, MBD3, MYNN, PRDM14, SIX5,* and *ZNF207*).

We labeled as “Repressed” the ten transcription factors with an NES of < 0 in the IDF, IDM, and SDID comparisons. These transcription factors are predicted to be repressed in female and male AD patients, with a significantly greater expression in female AD patients than in male AD patients (Genes - *GTF2B, HOXB13, NANOG, NME2, PCGF2, SNAI2, ZBTB7A, ZC3H8, ZHX1,* and *ZHX2*).

The two “Divergent” transcription factors possessed NES values with different signs in the IDF, IDM, and SDID comparisons. In the case of *CEBPZ*, results indicate activation in female AD patients and repression in male AD patients; meanwhile, results indicate *TERF2* activation in male AD patients and repressed female AD patients.

## DISCUSSION

AD patients suffer from progressive memory loss and other cognitive features, although etiology and pathogenesis remain complex. Recent omics-based studies have focused on exploring the influence of sex on neurodegenerative disease [54, 55]. Although discrepancies exist, a large collective of studies has provided evidence for the influence of sex on risk factors, prevalence and incidence, clinical manifestation, and treatment response [8, 56, 57]; therefore, evaluating the impact of sex-based differences on AD remains critical to improving prevention, diagnosis, and outcomes in female and male patients. We investigated sex-based differences in the molecular mechanisms associated with AD by performing two specific meta-analyses in brain regions primarily impacted by AD – the CT and HP. While the CT represents the central region involved in cognitive functions, the HP takes part in mental processes related to memory and those related to the production and regulation of emotional states and spatial navigation.

Our results demonstrated that gene expression in female AD patients is more significantly affected than in male AD patients. We found no significantly differentially expressed genes in the SDID comparison in the HP; however, analysis of the CT revealed 389 significantly differentially expressed genes in the SDID comparison. Therefore, this work focused on CT and the genes repressed in female AD patients (Subset I) that significantly impacted the functional profiling. Consistent with our findings, a multi-omics review performed by Lei Guo and colleagues stated that only 20% of selected omics studies of sex-based differences in AD describe male-specific changes, with molecular-level changes primarily described in females [57]. We detected significantly differentially expressed genes in the SDID comparison in female AD patients in the CT, the central region of the brain involved in cognitive function. This may represent a potential reason females present with a more rapid decline and cognitive deterioration [58].

We divided the resulting network of the Subset I genes in female AD patients from the SDID comparison into 10 clusters whose significantly altered processes mainly affected signal neurotransmission, Aβ aggregation, and deposition, which all critically impact cognitive impairment. Wang et al. reported the *ATP6V1A* gene in cluster 2 as a critical regulator in AD and related to cognitive deficits using a multi-omics approach that was later experimentally confirmed [59]. The downregulation of these critical genes in females may contribute significantly to the sex-based differences associated with AD.

Female AD patients also displayed dysregulation in genes encoding glutamate and GABA neurotransmitters, which would disturb the synaptic circuit (presynaptic, postsynaptic, and transmission). Synaptic loss significantly correlates with cognitive deficits in AD [60]; furthermore, neurotransmitter imbalance can influence synaptic plasticity (the fundamental ability of synapses to change their strength), which plays a crucial role in learning and memory [61]. Glutamate - the most important excitatory neurotransmitter in the central nervous system with particular relevance in memory and recovery - is also considered the primary mediator of sensory, motor, cognitive, and emotional information [62]. Conversely, GABA, which is widely distributed in the neurons of the CT, acts as an inhibitory messenger, thereby stopping the action of excitatory neurotransmitters. Additionally, GABA contributes to motor control, vision, and anxiety regulation, among other CT functions [63].

The interaction between L1 and Ankyrin proteins also appeared significantly altered in female AD patients. Several components of this family of scaffold proteins have been linked to neurodegenerative diseases (including AD) through the modulation of neuronal excitability and neuronal connectivity via ion channels [64, 65].

The accumulation of misfolded proteins in the human brain represents a critical factor in many neurodegenerative diseases, including AD [66]; indeed, a balance between the mechanisms that mediate protein folding, elimination, trafficking, Aβ aggregation, and deposition remains crucial to AD development. Female AD patients present with dysregulated pH values and calcium-mediated signaling, which are integrally involved in synapsis and neurotoxicity [67]. Aging is associated with more significant brain acidosis, which may affect AD-associated pathophysiological processes such as Aβ aggregation or inflammation [68]. Meanwhile, calcium signaling can promote the accumulation of Aβ plaques and neurofibrillary tangles in the brain [69] and remains critical for synaptic activity and memory, a process closely related to signal transduction pathways such as the MAPK kinase pathway [70], which also undergoes significant alterations in female AD patients. Said patients also presented alterations in the amino acids that regulate mTORC1 signaling, which relates to autophagy inhibition, clearance, and abnormal protein formation in human neurons and mice [71, 72]. Sex-based differences in autophagy and the association with AD have been extensively reviewed [73], with the authors proposing that lower basal autophagic activity observed in females leads to lower levels of neural protection mediated by the clearance of aggregated proteins such as Aβ and hyper-phosphorylated Tau tangles. This mechanism would increase the risk of AD and greater pathologic severity in females. Furthermore, autophagy requires the suppressed transcription of genes such as *ATP6AP2* [74], which we observed in female AD patients in this study.

Protein folding and the HSP90 chaperone cycle for steroid hormone receptors represent additional functions related to protein abnormalities that displayed alterations in female AD patients. The steroid receptor-mediated regulation of HSP90 function remains critical for their signal-transducing function and their participation in the folding, stabilization, and trafficking of proteins [75]. Imbalances affecting the function of HSP90 and other chaperones can contribute to tauopathies, AD onset, and disease progression [76, 77]; moreover, imbalances/abnormalities in endocytotic processes represent among the earliest alterations associated with AD. The dysregulation of clathrin-mediated endocytosis in female AD patients plays a significant role in the internalization of amyloid precursor protein (APP) and Aβ generation [78]. Aβ generation and cognitive decline also is associated with NFAT activation [79], which we found altered and related to dectin-1 in female AD patients. COP9 signalosome assembly participates in ubiquitin-mediated proteolysis [80], and the disruption of COP9 signalosome in parasites causes the dysregulation of the ubiquitin-proteasome pathway, therefore impacting protein degradation and cell death [81]. The dysregulation of Protein K11-linked ubiquitination, which functions in protein degradation and inflammation, also represents an essential factor [82–84].

We discovered 23 transcription factors with significant differential expression in the SDID comparison, which we then classified into three groups considering their expression profile in the IDF and IDM comparisons. While 11 transcription factors displayed an “activated” profile in female and male AD patients, the elevated levels observed in females make these transcription factors potential targets. Studies have linked aberrant ADNP expression to neural developmental disorders and proposed as a novel marker for the onset of frontotemporal dementia [85, 86]. Seefelder et al. linked HMN3 to Huntington’s disease [87], while IRF3 participates in AD progression and cognitive impairment [88]. Studies have linked KFL5 and KFL9 to AD progression and development [89, 90], MAZ participates in amyloidosis [91], MBD3 induces neurotoxicity in mice [92], and MYNN has significant diagnostic value in AD patients [93]. Studies have linked PRDM14 to pluripotency, motor neurons, and brain germinoma [94], while Seznec et al. associated SIX5 with abnormal tau expression [95].

We also found several “Repressed” transcription factors in female and male AD patients but with significantly greater repression in females. Lee et al. identified GTF2B as a dysregulated transcription factor in AD [96]; however, HOXB13, previously linked to prostate cancer risk [97, 98], has been linked with AD for the first time in this study. Nanog is a pluripotency stem cell marker important in stem cell therapies for neurodegenerative diseases [99, 100]. Studies have linked NME proteins in cancer progression [101]; however, this study represents the first link between NME2 in AD and any neural disorders. Finally, Florentinus-Mefailoski et al. reported increased levels of PCFG2 in AD [102]. Our study also linked multiple zinc-finger transcription factors related to cancer progression and neural development to AD for the first time. The EMT-transcription factor SNAI2 is vital for neural crest specification, cell migration, and survival during neural crest development and has therapeutic implications in neuroblastoma cells [103]. Studies in hepatocarcinoma cells reported that ZBTB7A may regulate neural development [104], while Zou et al. described Zc3h8 as a repressor of inflammation in zebrafish [105]. ZHX1 and ZHX2 participate in the development and progression of several types of cancer, and the low expression of both impacts a poor prognosis in chronic lymphocytic leukemia [106].

CEBPZ and TERF2 represent transcription factors with divergent gene expression patterns in females and males. CEBPZ, which belongs to a family of transcription factors relevant to immune response control and inflammation [107], is “Activated” in female AD patients and “Repressed” in male AD patients. Studies have associated additional components of this family with neuroinflammation, an early event in AD [108, 109]. TERF2 is “Repressed” in female AD patients and “Activated” in male AD patients and has been related to senescence in AD [110, 111]. Wu et al. positively correlated the expression of TRF1 and TRF2 in AD patients with age and Tau protein levels in blood serum [112].

### Strengths and limitations

Exploring sexual bias may significantly improve clinical outcomes in female and male AD patients; we employed an *in-silico* approach using computational models as powerful tools for evaluating and integrating data. As the sample size increases with the number of studies integrated into the meta-analysis, we can detect more subtle effects and provide greater consensus and statistical power in the obtained results [54, 55, 113–115]. Additionally, our *in-silico* analysis is based on FAIR data (Findable, Accessible, Interoperable, Reusable) [116], which we believe to be particularly relevant; indeed, we believe the sharing and reusing of research data to be critical in making advances. *In-silico* integrative approaches to analyze sex-based differences in gene transcription in AD patients have been carried out; for example, Paranjpe et al. systematically meta-analyzed RNA-sequencing data collected from brain samples [117]. Instead, our study discriminates between the two primarily affected brain regions (CT and HP) in two different meta-analyses, leading to more region-specific results. Moreover, we conducted our analyses using the limma R package for individual differential expression analyses [26] and then the metafor R package for the gene expression meta-analyses [29], which allowed us to analyze data following our SDID comparison, which includes four experimental groups and identifies sex-based differences in AD, considering the inherent variability among males and females in healthy conditions. The joint use of the IDM, IDF, and SDID comparisons allowed us to more precisely classify the genes with significantly altered expression levels in the significant comparison into ten clusters according to their expression profile across the four experimental groups.

Our study has some noteworthy limitations. First, while our methodology favors the detection of robust expression patterns across the different studies included, it may also mask more subtle expression patterns specific to the different subtypes of AD. In this regard, due to the lack of information in some studies, the type of AD (LOAD and EOAD) or the genotypic status of the APOE gene has not been included as covariates in the analysis.Our study has some noteworthy limitations. First, while our methodology favors the detection of robust expression patterns across the different studies included, it may also mask more subtle expression patterns specific to the different subtypes of AD. In this regard, due to the lack of information in some studies, the type of AD (LOAD and EOAD) or the genotypic status of the APOE gene has not been included as covariates in the analysis.

Lastly, cumulative evidence points to the influence of sex in various disorders; however, it remains challenging to encounter data segregation by sex in research studies, even in those conducted to explore diagnostic/prognostic factors. For example, we excluded 40 (48%) studies from our systematic review due to the absence of information regarding sex; therefore, we highlight the need to include information regarding sex in research studies and databases, given their vast relevance to health.

### Perspectives and significance

The results obtained in this study contribute to a better understanding of the impact of sex-based differences at molecular and functional levels in AD. The genes, biological processes, pathways, and transcription factors identified in the SDID comparison represent a source of valid targets to improve clinical outcomes and the springboard for future research as comparisons with other studies.

## CONCLUSIONS

Our results highlight sex-based differences as more evident in female AD patients, particularly affecting the cerebral cortex whose neurons are mainly responsible for cognitive processing. We identified impairments in the primary neurotransmitters (glutamate and GABA) and different mechanisms for generating/depositing Aβ plaques. We also identified genes and transcription factors representing novel options that may guide new therapeutic strategies. More studies that consider sex as a critical dimension are required for a better interpretation of the results and to avoid masking subtle differences that could impact tailored interventions. Lastly, we underline the relevance of sharing data and working in open platforms for scientific progress.

## Declarations

### Ethics approval and consent to participate

Not applicable

## Consent for publication

Not applicable

## Availability of data and materials

The data used for the analyses described in this work are publicly available at GEO (16). The accession numbers of the GEO datasets downloaded are GSE118553, GSE125583, GSE1297, GSE132903, GSE15222, GSE29378, GSE37263, GSE48350, GSE5281, and GSE84422.

## Competing interests

The authors declare no competing interests

## Funding

This research was supported by and partially funded by the Institute of Health Carlos III (project IMPaCT-Data, exp. IMP/00019), co-funded by the European Union, European Regional Development Fund (ERDF, “A way to make Europe”), and PID2021-124430OA-I00 funded by MCIN/AEI/10.13039/501100011033/ FEDER, UE (”A way to make Europe”). Irene Soler-Sáez thanks the Spanish Ministry of Universities for her predoctoral grant FPU20/03544.

## Authors contributions

ALC analyzed the data; FGG designed and supervised the bioinformatics analysis; ISS, and ALC designed and implemented the web tool; ALC, ZA, ISS, AM, FRG, and FGG wrote the manuscript; ALC, ZA, ISS, AM, FRG, MRH, AP, MPM, SL, and FGG helped in the interpretation of the results; all authors writing-review and editing; FGG conceived the work. All authors read and approved the final manuscript.

## Acknowledgments

The authors thank the Principe Felipe Research Center (CIPF) for providing access to the cluster, co-funded by European Regional Development Funds (FEDER) in the Valencian Community 2014-2020. The authors also thank Stuart P. Atkinson for reviewing the manuscript.

## Notes

### Competing Interest Statement

The authors have declared no competing interest.

### Summary of Updates

Improvement writing manuscript

